# Gene regulatory network architecture in different developmental contexts influences the genetic basis of morphological evolution

**DOI:** 10.1101/219337

**Authors:** Sebastian Kittelmann, Alexandra D. Buffry, Franziska A. Franke, Isabel Almudi, Marianne Yoth, Gonzalo Sabaris, Juan Pablo Couso, Maria D. S. Nunes, Nicolás Frankel, José Luis Gómez-Skarmeta, Jose Pueyo-Marques, Saad Arif, Alistair P. McGregor

## Abstract

Convergent phenotypic evolution is often caused by recurrent changes at particular nodes in the underlying gene regulatory networks (GRNs). The genes at such evolutionary ‘hotspots’ are thought to maximally affect the phenotype with minimal pleiotropic consequences. This has led to the suggestion that if a GRN is understood in sufficient detail, the path of evolution may be predictable. The repeated evolutionary loss of larval trichomes among *Drosophila* species is caused by the loss of *shavenbaby* (*svb*) expression. *svb* is also required for development of leg trichomes, but the evolutionary gain of trichomes in the ‘naked valley’ on T2 femurs in *Drosophila melanogaster* is caused by the loss of *microRNA-92a* (*miR-92a*) expression rather than changes in *svb*. We compared the expression and function of components between the larval and leg trichome GRNs to investigate why the genetic basis of trichome pattern evolution differs in these developmen*tal* contexts. We found key differences between the two networks in both the genes employed, and in the regulation and function of common genes. These differences in the GRNs reveal why mutations in *svb* are unlikely to contribute to leg trichome evolution and how instead *miR-92a* represents the key evolutionary switch in this context. Our work shows that variability in GRNs across different developmen*tal* contexts, as well as whether a morphological feature is lost versus gained, influence the nodes at which a GRN evolves to cause morphological change. Therefore, our findings have important implications for understanding the pathways and predictability of evolution.

**Author Summary:** A major goal of biology is to identify the genetic cause of organismal diversity. Convergent evolution of traits is often caused by changes in the same genes – evolutionary ‘hotspots’. *shavenbaby* is a ‘hotspot’ for larval trichome loss in *Drosophila*, however *microRNA-92a* underlies the gain of leg trichomes. To understand this difference in the genetics of phenotypic evolution, we compared the expression and function of genes in the underlying regulatory networks. We found that the pathway of evolution is influenced by differences in gene regulatory network architecture in different developmen*tal* contexts, as well as by whether a trait is lost or gained. Therefore, hotspots in one context may not readily evolve in a different context. This has important implications for understanding the genetic basis of phenotypic change and the predictability of evolution.

## Introduction

A major challenge in biology is to understand the relationship between genotype and phenotype, and how genetic changes modify development to generate phenotypic diversification. The genetic basis of many phenotypic differences within and among species have been identified [e.g. 1,2-15], and these findings support the generally accepted hypothesis that morphological evolution is predominantly caused by mutations affecting cis-regulatory modules of developmen*tal* genes [16]. Moreover, it has been found that changes in the same genes commonly underlie the convergent evolution of traits [reviewed in 17]. This suggests that there are evolutionary ‘hotspots’ in GRNs: changes at particular nodes are repeatedly used during evolution because of the role and position of the gene in the GRN, and the limited pleiotropic effect of the change [18-21].

The regulation of trichome patterning is an excellent system for studying the genetic basis of morphological evolution [22]. Trichomes are actin protrusions from epidermal cells that are overlaid by cuticle and form short, non-sensory, hair-like structures. They can be found on various parts of insect bodies during different life stages, and are thought to be involved in, for example, thermo-regulation, aerodynamics, oxygen retention in semi-aquatic insects, grooming, and larval locomotion [23-27] (Fig 1).

**Fig 1.**
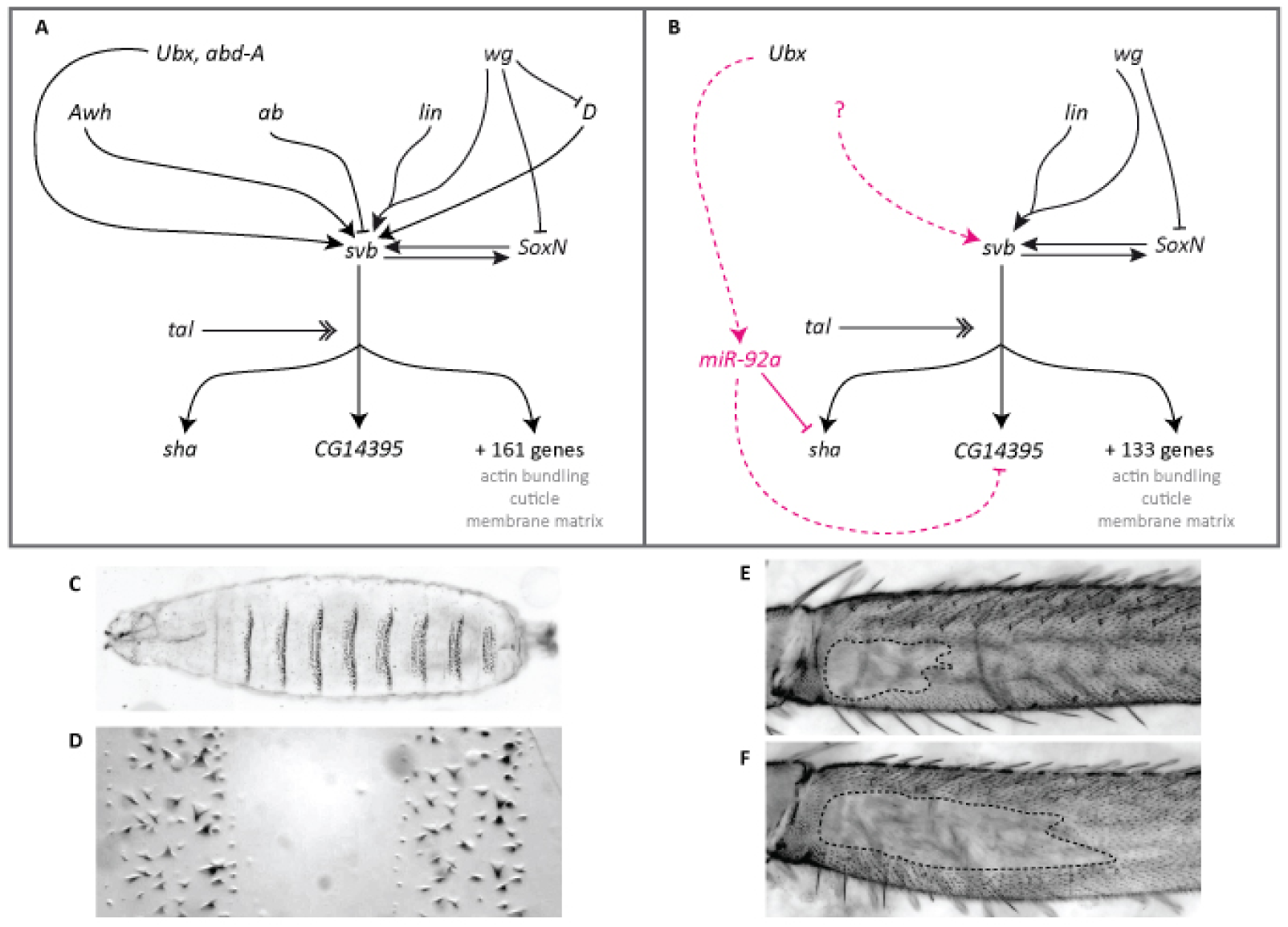
The GRNs controlling formation of trichomes on larval and leg epidermis differs between these developmen*tal* contexts. (A) Simplified GRN for larval trichome development [22,29,75,76]. (B) GRN for leg trichome development. Magenta colour indicates interactions found only during leg development. Dotted lines indicate likely interactions. Expression of *svb* is controlled by several upstream transcription factors and signalling pathways some of which are not active during leg trichome development. The question mark indicates that there are likely to be other unknown activators of *svb* in legs. Activation of Svb protein requires proteolytic cleavage by small peptides encoded by *tal* [14,31,77]. Active Svb then regulates the expression of at least 161 target genes in embryos, the expression of 133 of which is detectable in legs [29,32]. The products of these downstream genes are involved in actin bundling, cuticle segregation, or changes to the matrix, which lead to the actual formation of trichomes. SoxN and Svb activate each other and act partially redundantly on downstream targets in larvae and it is hypothesise that this interaction also occurs in legs based on expression data [33,35]. *miR-92a* is only expressed in naked leg cells where it represses *sha* and possibly *CG14395* and thereby acts as a short circuit for *svb*. Its expression is likely controlled by *Ubx*. (C, D) Trichomes on the ventral side of the larval cuticle form stereotypic bands (‘denticle belts’) separated by trichome-free cuticle. (E, F) A trichome-free region on the posterior of the T2 femur differs in size between different strains. Shown are *OregonR* (E) and e^4^, wo^2^, ro^2^ (F).

The GRN underlying trichome formation on the larval cuticle of *Drosophila* species has been characterised in great detail [reviewed in 21,22,28] (Fig 1). Several upstream transcription factors, signalling pathways, and *tarsal-less* (*tal*)-mediated post-translational proteolytic processing lead to the activation of the key regulatory transcription factor Shavenvbaby (Svb), which, with SoxNeuro (SoxN), activates a battery of downstream effector genes [14,29-35]. These downstream factors modulate cell *sha*pe changes, actin polymerisation, or cuticle segregation, which underlie the actual formation of trichomes [29,32]. Importantly, ectopic activation of *svb* during embryogenesis is sufficient to drive trichome development on otherwise naked larval cuticle, and loss of *svb* function leads to a loss of larval trichomes [36].

Regions of dorso-lateral larval trichomes have been independently lost at least four times among *Drosophila* species [37,38]. In all cases, recombination mapping and functional analyses have shown that this phenotypic change is caused by changes in several *svb* enhancers, resulting in a loss of *svb* expression [6,9,37-40]. The modular enhancers of *svb* are thought to allow the accumulation of mutations that facilitate the loss of certain larval trichomes without deleterious pleiotropic consequences. It is thought that evolutionary changes in larval trichome patterns cannot be achieved by mutations in genes upstream of *svb* because of deleterious pleiotropic effects, while changes in individual *svb* target genes would only affect trichome morphology rather than their presence or absence [19-21,29,32]. Given the position and function of *svb* in the larval trichome GRN, these data suggest that *svb* is a hotspot for the evolution of trichome patterns more generally because it is also required for the formation of trichomes on adult epidermis and can induce ectopic trichomes on wings when over expressed [36,41]. Therefore, one could predict that changes in adult trichome patterns are similarly achieved through changes in *svb* enhancers [20,21].

The trichome pattern on femurs of second legs also varies within and between *Drosophila* species [1,42] (Fig 1). In *D. melanogaster*, an area of trichome-free cuticle or ‘naked valley’ varies in size among strains from small to larger naked regions. Other species of the *D. melanogaster* species subgroup only exhibit larger naked valleys [1,42]. Therefore, trichomes have been gained at the expense of naked cuticle in some strains of *D. melanogaster*. Differences in naked valley size between species have been associated with differences in the expression of *Ultrabithorax* (*Ubx*), which represses the formation of leg trichomes [42]. However, genetic mapping experiments and expression analysis have shown that naked valley size variation among populations of *D. melanogaster* is caused by cis-regulatory changes in *miR-92a* [1]. This microRNA represses trichome formation by repressing the *svb* target gene *sha*venoid (*sha*), and *D. melanogaster* strains with small and large naked valleys exhibit weaker or stronger *miR-92a* expression, respectively, in developing femurs [1,43]. Therefore, while *svb* is thought to be a hotspot for the evolutionary loss of patches of larval trichomes, it does not appear to underlie the evolutionary gain of leg trichomes in *D. melanogaster*.

Differences in GRN architecture among developmen*tal* contexts may affect which nodes can evolve to facilitate phenotypic change in different tissues or developmen*tal* stages. In addition, an evolutionary gain or loss of a phenotype may also result from changes at different nodes in the underlying GRN, i.e. alteration of a particular gene may allow the loss of a trait but changes in the same gene may not necessarily result in the gain of the same trait. Therefore, a better understanding of the genetic basis of phenotypic change and evaluation of the predictability of evolution requires characterising the expression and function of GRN components in different developmen*tal* contexts and studying how the loss versus the gain of a trait is achieved.

Here we report our comparison of the regulation of trichome development in legs versus embryos. Our results show differences in expression and function of key components of the GRN between these two developmen*tal* contexts. These differences indicate that *svb* is likely unable to act as a switch for the gain of leg trichomes because it is already expressed throughout the legs in both naked and trichome-producing cells. Instead, regulation of *sha* by *miR-92a* appears to act as the switch between naked and trichome-producing cells in the leg. This shows that differences in GRNs between different developmen*tal* contexts can affect the pathway used by evolution to generate phenotypic change.

## Results

### Differences between genes expressed during leg and larval trichome development

To better characterise the regulation of leg trichome development we first carried out RNA-Seq of T2 pupal legs between 20 and 28 hours after puparium formation (hAPF): the window when leg trichomes are specified [42] (S1-6 Files). We found that key genes known to be involved in larval trichome formation are expressed in legs. These include *Ubx, SoxN, tal, svb*, and *sha*, as well as key components of the Delta-Notch, Wnt and EGF signalling pathways (Fig. 1, S1 Fig and S1 Table). However, expression of several genes known to regulate larval trichome development [29,32,35] is barely detectable in legs (i.e. below or around 1 FPKM). These include Dichaete, *Arrowhead*, and *abrupt*, which are also known to regulate *svb* expression during larval trichome development [33,40] (Fig. 1 and S1 Table). Furthermore, the expression of 28 of the 163 known targets of *svb* in embryos [29,32] are barely detectable in our dataset (FPKM at or below 1) (S2 Table). In addition, 10 out of the 43 genes thought to be involved in larval trichome formation independently of *svb* [32,35] are also expressed at levels less than 1 FPKM in legs (S3 Table). Therefore, our RNA-Seq data shows substantial differences in both upstream and downstream components of the leg trichome GRN when comparing it to what is known for the embryonic GRN that specifies larval trichomes.

Our leg RNA-Seq data also allowed us to compare expression between strains of *D. melanogaster* with different sizes of naked valley: Oregon R (OreR) which has a small naked valley and *ebony* ^4^, *white ocelli* ^1^, rough^1^ (eworo) which has a large naked valley (Fig. 1). The size of the naked valley in these two strains is caused by differential expression of *miR-92a* [1]. We found that none of the known regulators of *svb* are differentially expressed between these two strains. In addition, we did not detect any significant differences in the expression of *svb* itself or most of its target genes including *sha* (S1 and S2 Tables). However, we found that *jing* interacting gene regulatory 1 (*jigr1*) is more highly expressed in the large naked valley strain eworo (S1 Table). *miR-92a* is usually co-expressed with this gene [44] since it is located in one of its introns, and therefore higher expression of *miR-92a* may be indirectly detectable in eworo. Note, however, while this expression difference is not significant after p value correction for false discovery rate (FDR), but that there is a clear trend towards higher expression of *jigr1* in eworo (S1 Table). These results are consistent with *miR-92a* mediated post-transcriptional regulation causing differences in naked valley size, and since this only occurs in a small proportion of leg cells, the effect on transcripts is likely to be difficult to detect using RNA-Seq.

### miR-92a is sufficient to repress leg trichomes and acts downstream of Ubx

We next sought to further examine the function of specific genes during leg trichome development compared to their roles in the formation of larval trichomes. It was previously shown that mutants of *miR-92a* have small naked valleys [45], which is consistent with the evolution of this locus underlying natural variation in naked valley size [1]. We confirmed these findings using a double mutant for *miR-92a* and its paralogue miR-92b [44], which exhibits an even smaller naked valley (Fig. 2). Note however, that we did not detect any changes to the larval trichome pattern in these mutants compared to heterozygotes. We examined the morphology of the proximal leg trichomes gained from the loss of *miR-92a* compared to the trichomes found more dis*tal*ly. We found that the trichomes gained were indistingui*sha*ble from the other leg trichomes (S1 Fig). This suggests that all of the genes required to generate leg trichomes are already transcribed in naked valley cells, but that *miR-92a* must be sufficient to block their translation. Indeed, we found that the extra trichomes that develop in the naked valley in the absence of *miR-92a* are dependent on *svb* because in a *svb* mutant background no trichomes are gained after loss of *miR-92a* (Fig 2). Furthermore, these results also show that trichome repression by *Ubx* in the naked valley [42] requires *miR-92a* because trichomes in the *miR-92a* mutant develop in the region where *Ubx* is expressed. Thus, our data shows that *Ubx* plays opposite roles in the larval and leg trichome GRNs: in embryos *Ubx* activates *svb* to generate larval trichomes [46], while we show that *Ubx*-mediated repression of leg trichomes [42,47] depends on *miR-92a* (Fig 1).

**Fig 2.**
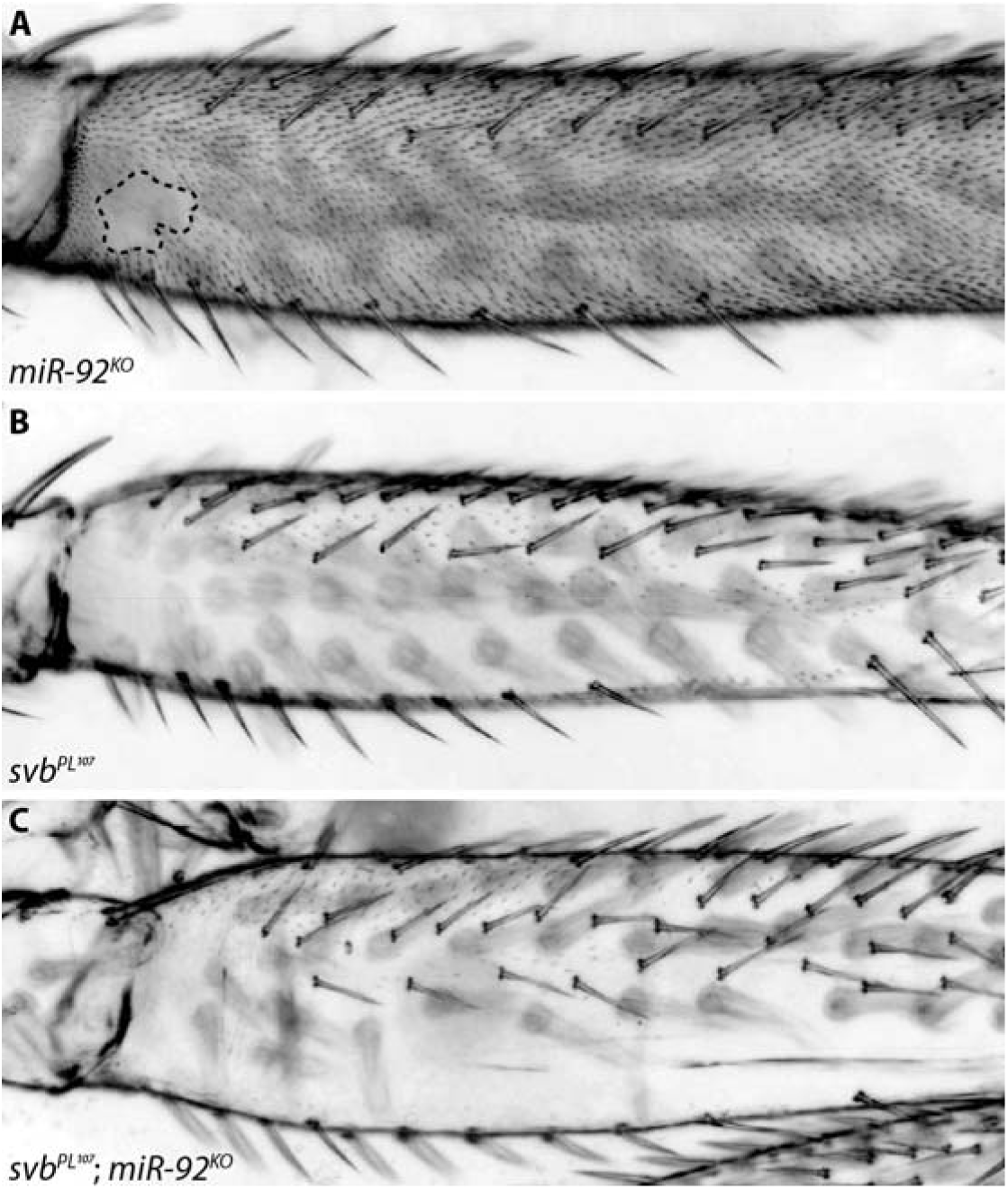
Leg trichome patterns in miR-92a/miR-92b mutants. (A) Flies mutant for both *miR-92a* and miR-92b gain trichomes in the naked valley. (B) Most trichomes on the posterior T2 femur are repressed in *svb*^*pL107*^ flies. (C) No trichomes are gained upon loss of *miR-92a* and miR-92b in a *svb* ^*pL107*^ background.

### Regulation of *svb* during leg trichome patterning

The results above suggest that *svb* is expressed in the naked valley but is unable to induce the formation of trichomes because of the presence of miR-92a. To test this further we examined the expression of *svb* transcripts in pupal T2 legs using in situ hybridization. However, this method produced inconsistent results among legs and it was difficult to distinguish between signal and background in the femur. Therefore we examined the expression of a nuclear GFP inserted into a BAC containing the entire *svb* cis-regulatory region, which was previously shown to reliably capture the expression of this gene [41]. We detected GFP throughout T2 legs at 24 hAPF including in the proximal region of the posterior femur (S2 Fig). This indicates that *svb* is expressed in naked valley cells that do not produce trichomes as well as in more dis*tal* trichome producing cells.

We next investigated the regulatory sequences responsible for *svb* expression in T2 legs. To do this we carried out ATAC-Seq [45,46] on chromatin from T2 legs during the window of 20 to 28 hAPF when leg trichomes are specified [42]. Embryonic expression of *svb* underlying larval trichomes is regulated by several enhancers spanning a region of approximately 90 kb upstream of the transcription start site of this gene [5,9] (Fig 3). Several of these larval enhancers also drive reporter gene expression during pupal development [41]. We observed that the embryonic enhancers DG3, E and 7 contained regions of open chromatin according to our T2 leg ATAC-Seq data. However, we found additional accessible chromatin regions that do not overlap with known *svb* embryonic enhancers (Fig 3).

**Fig 3.**
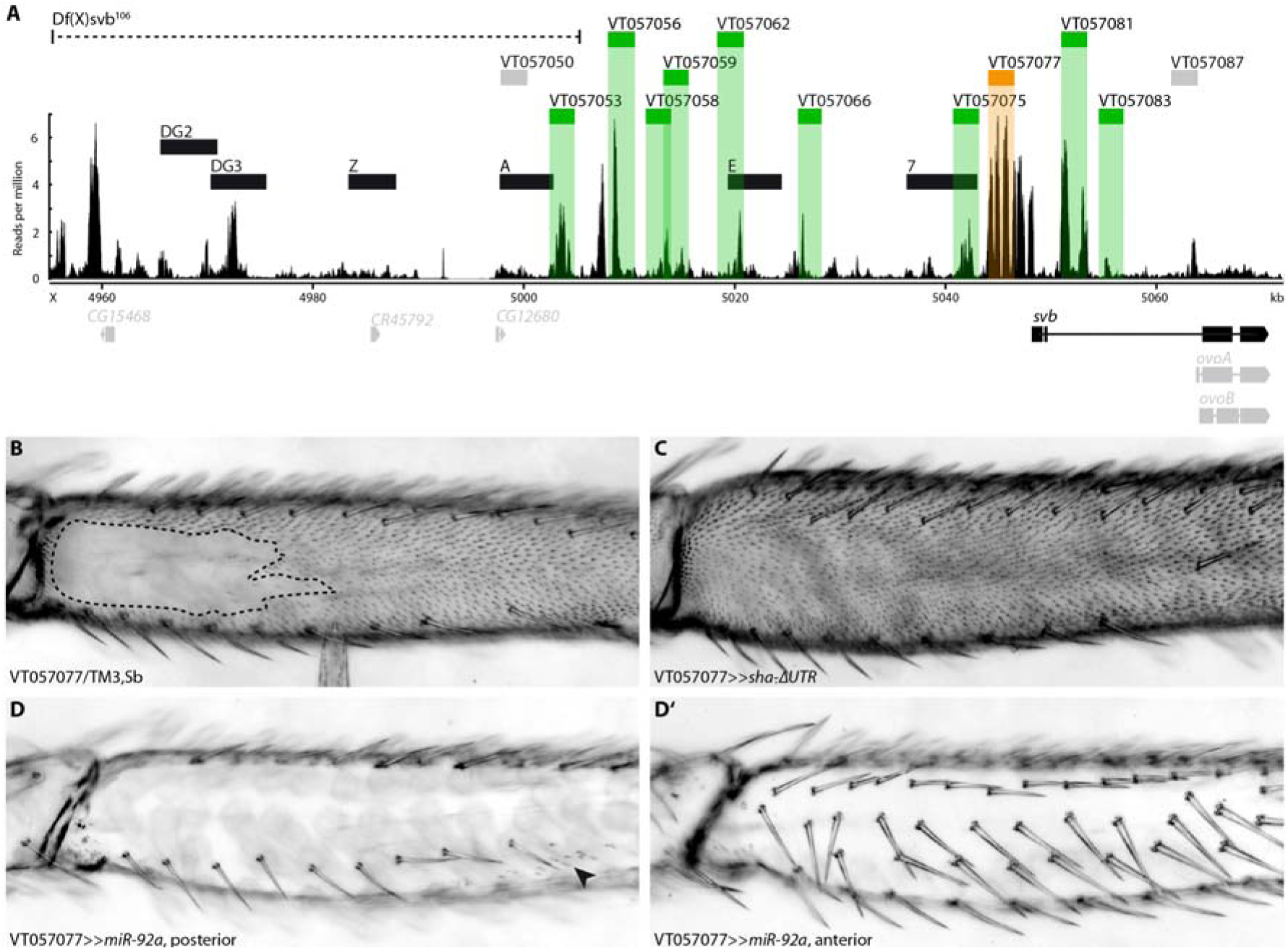
Enhancers of *svb*. (A) Overview of the chromatin accessibility profile (ATAC-seq) at the *ovo*/*svb* locus. Indicated are: the deficiency used (dotted line), known larval *svb* enhancers (black boxes), and tested putative enhancers (grey boxes: no expression in pupal legs, green/orange boxes: expression in pupal legs). Region VT057077 (orange) is able to drive expression during trichome formation (see B-D). The bottom panel shows expressed variants of genes at the locus (black) and genes/variants not expressed (grey). Boxes represent exons, lines represent introns. (B) VT057077 has a naked valley of intermediate size. (C) Expression of *sha*-ΔUTR under its control induces trichome formation in the naked valley. (D, D’) Driving *miR-92a* with VT057077 represses trichome formation on the anterior and posterior of the second leg femur. Small patches of trichomes can sometimes still be found (arrowhead).

Deletion of a region including the embryonic enhancers DG2 and DG3 [Df(X)svb^108^] (Fig 3) results in a reduction in the number of dorso-lateral larval trichomes when in a sensitized genetic background or at extreme temperatures [5]. Moreover, Preger-Ben Noon and colleagues (2017) [41] recently showed that this deletion, as well as a larger deletion that also removes embryonic enhancer A ([Df(X)*svb*^106^], see Fig 3), results in the loss of trichomes on abdominal segment A5, specifically in males. We found several peaks of open chromatin in the regions covered by these two deficiencies in our second leg ATAC-seq dataset (Fig 3) and therefore tested the effect of *Df*(*X*)*svb*106 on leg trichome development. We found that deletion of this region and consequently enhancers DG2, DG3, Z and A did not affect the size of the naked valley or the density of trichomes on the femur or other leg segments of flies raised at 17°C, 25°C, or 29°C (compared to the paren*tal* lines) (S3 Fig). This suggests that while this region may contribute to *svb* expression in legs, its removal does not perturb the robustness of leg trichome patterning.

Next, to try to identify enhancer(s) responsible for leg expression, we employed all available GAL4 reporter lines for cis-regulatory regions of *svb* (S4 Table) that overlap with regions of open chromatin downstream of the above deficiencies (Fig 3). All 10 regions that overlap with open chromatin are able to drive GFP expression to some extent in second legs between 20 and 28 hAPF, as well as in other pupal tissues (S4 Fig). While some of the regions only produce expression in a handful of epidermal cells or particular regions of the T2 legs, none are specific to the presumptive naked valley. Moreover, VT057066, VT057077, VT057081, and VT057083 appear to drive variable levels of GFP expression throughout the leg (S4 Fig). Note that the two regions overlapping with larval enhancers E and 7 (VT057062 and VT057075, respectively) only drive weak expression in a few cells in the tibia and tarsus (S4 Fig).

To further test whether the expression of any of these regions is consistent with a role in trichome formation, we used them to drive expression of the trichome repressor *miR-92a* and the trichome activator *sha*-ΔUTR [see 1]. Intriguingly, driving *miR-92a* under control of only one of the fragments (VT057077) caused the repression of trichomes on all legs (Fig 3 and S5 Fig) as well as on wings and halteres (S5 Fig). Expressing *miR-92a* under control of VT057062 or VT057075 had no noticeable effect, and with two of the other fragments (VT057053, VT057056) only led to repression of trichomes in small patches along the legs consistent with the GFP expression pattern (S4 and S5 Figs).

Driving *sha*-ΔUTR with VT057077 is sufficient to induce trichome formation in the naked valley (Fig 3) and on the posterior T3 femur (S5 Fig). Driving *sha*-ΔUTR under control of any of the other nine regions did not produce any ectopic trichomes in the naked valley on T2 or on any other legs. These results indicate that a single enhancer, VT057077, is able to drive *svb* expression throughout the second leg in both regions which normally produce trichomes and in naked areas.

### svb and *sha* differ in their capacities to induce trichomes in larvae and legs

It was previously shown that *miR-92a* inhibits leg trichome formation by repressing translation of the *svb* target *sha* [1]. However, *sha* mutants are still able to develop trichomes in larvae albeit with abnormal morphology [29]. These data suggest that there are differences in the functionality of *svb* and *sha* in larvae versus leg trichome formation, and therefore we next verified and tested the capacity of *svb* and *sha* to produce larval and leg trichomes.

As previously shown [29], ectopic expression of *svb* is sufficient to induce trichome formation on normally naked larval cuticle (Fig 4). However, we found that ectopic expression of *sha* in the same cells does not lead to the production of trichomes (Fig 4). *svb* is also required for posterior leg trichome production [41] (Fig 2 and S6 Fig), but over expression of *svb* in the naked valley does not produce ectopic trichomes (Fig 4). Over expression of *sha* on the other hand is sufficient to induce trichome development in the naked valley [1] (Fig 4). These results show that *svb* and *sha* differ in their capacities to generate trichomes in larvae versus legs.

**Fig 4.**
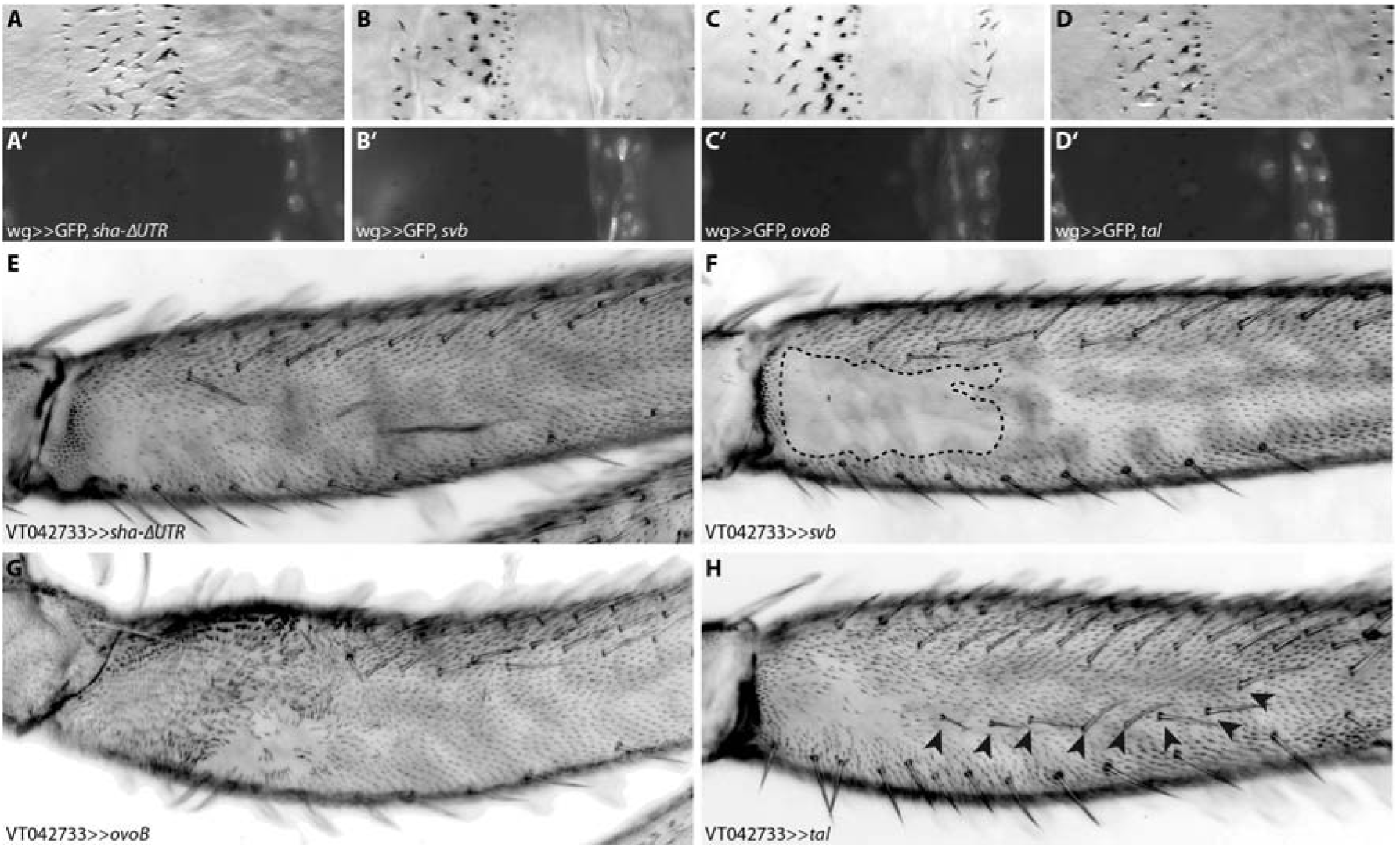
Ectopic trichome formation on naked cuticle. Driving *sha*-ΔUTR (A) under control of wg-Gal4 does not lead to ectopic trichome formation on otherwise naked larval cuticle. Driving *svb* (B) or its constitutively active variant *ovoB* (C) is sufficient to activate trichome development, but expressing only the Svb activator *tal* (D) is not. GFP was co-expressed in each case to indicate the wg expression domain (A’-D’). Ectopic activation of *sha*-ΔUTR in the proximal femur (E) is able to induce trichome formation, but ectopic *svb* (F) is not. Driving either *ovoB* (G) or the activator *tal* (H) leads to ectopic trichome development. Expression of *ovoB* has additional effects on leg development (e.g. a bending of the proximal femur), while expression of *tal* also leads to the development of ectopic bristles on the femur (arrowheads in H).

Interestingly, we observed that the ectopic trichomes produced by expression of *sha*-ΔUTR in the naked valley are significantly shorter than those on the rest of the leg (S1 Fig). This suggests that although *sha* is able to induce trichome formation in these cells, other genes are also required for their normal morphology. We observed that another characterised *svb*-target gene, *CG14395* [32], is also a strongly predicted target of miR-92a: its 3’UTR contains two conserved complete 8-mers corresponding to the binding site for this microRNA. We found that *CG14395* is also expressed in pupal second legs according to our leg RNA-Seq data (S2 Table) and furthermore RNAi against this gene resulted in shorter leg trichomes (S7 Fig) Therefore it appears that *miR-92a* also represses *CG14395* and potentially other *svb* target genes in addition to *sha* to block trichome formation.

### Over expression of *tal* or *ovoB* can induce trichomes

Svb acts as a transcriptional repressor and requires cleavage by the proteasome to become a transcriptional activator. This cleavage is induced by small proteins encoded by the *tal* locus [14,30,31]. We therefore tested if *svb* is unable to promote trichome development in the naked valley because it is not activated in these cells. We found that expressing the constitutively active form *ovoB*, or *tal*, in naked leg cells is sufficient to induce trichome formation (Fig 4), which is consistent with loss of trichomes in *tal* mutant clones of leg cells (S6 Fig). Furthermore, it appears that *tal*, like *svb*, is expressed throughout the leg (S6 Fig). It follows that *svb* and *tal* are expressed in naked cells but are unable to induce trichome formation under normal conditions because of repression of *sha, CG14395* and possibly other genes by miR-92a. We hypothesise that over expression of *tal* on the other hand must be able to produce enough active Svb to result in an increase of *sha* transcription to overwhelm *miR-92a* repression.

## Discussion

### The GRNs for larval and leg trichome patterning differ in composition and evolution

The causative genes and even nucleotide changes that underlie the evolution of an increasing number and range of phenotypic traits have been identified [17]. An important theme that has emerged from these studies is that the convergent evolution of traits is often explained by changes in the same genes – so called evolutionary ‘hotspots’ [17,48]. This suggests that the architecture of GRNs may influence or bias the genetic changes that underlie phenotypic changes [18,19,21]. However, relatively little is known about the genetic basis of changes in traits in different developmen*tal* contexts and when features are gained versus lost [18].

It was shown previously that changes in the enhancers of *svb* alone underlie the convergent evolution of the loss of larval trichomes, while the gain of leg trichomes in *D. melanogaster* is instead mainly explained by evolutionary change in cis-regulatory regions of *miR-92a* [1,6,9,37-39]. We investigated this further by comparing the GRNs involved in both developmen*tal* contexts and examining the regulation and function of key genes.

Our results show that there are differences between the GRNs underlying the formation of larval and leg trichomes in terms of the expression of components and their functionality. These changes are found both in upstream genes of the GRN that help to determine where trichomes are made and in downstream genes whose products are directly involved in trichome formation (Fig 1). The latter may also determine the differences in the fine-scale morphology of these structures on larval and leg cuticle (Fig 1)[29].

Furthermore, while the key evolutionary switch in embryos, the gene *svb*, is also necessary for trichome production on the posterior leg, this gene is not sufficient to produce leg trichomes in the naked proximal region of the T2 femur. This is because the leg trichome GRN employs miR-92a, which inhibits trichome production by blocking the translation of the *svb* target gene *sha* and probably other target genes including *CG14395*. In the legs of *D. melanogaster, miR-92a* therefore acts as the evolutionary switch for trichome production, and consequently the size of the naked valley depends on the expression of this gene (Fig 5) [1].

**Fig 5.**
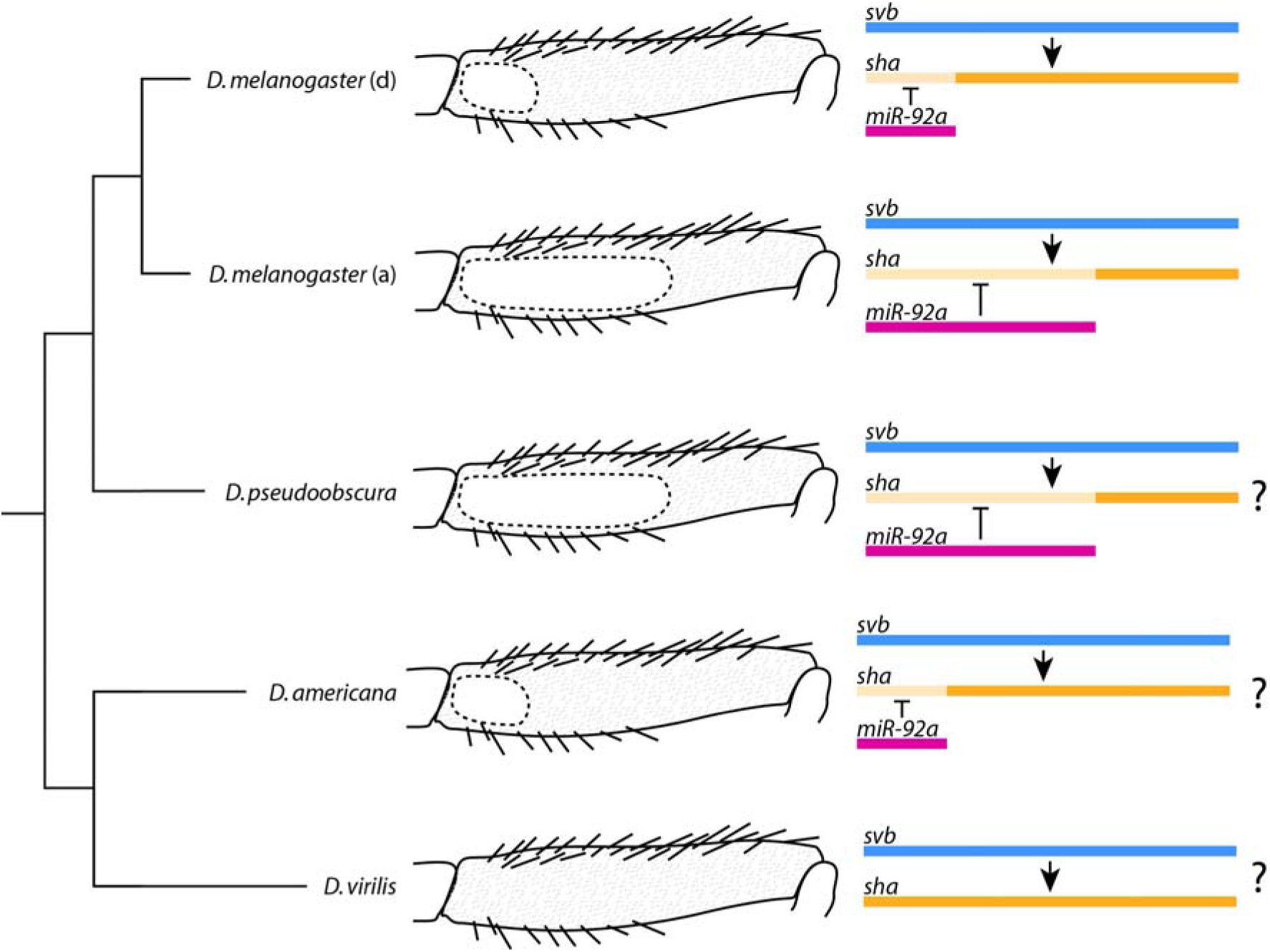
The size of the naked valley differs between and within species and is dependent on *miR-92a* expression. Loss of *miR-92a* expression in *D. melanogaster* has led to a derived (d) smaller naked valley in some populations while the ancestral state (a) is thought to be a large naked valley like in other melanogaster group species and other species (e.g. *D. pseudoobscura*). The absence of a naked valley in *D. virilis* is likely due to absence of *miR-92a* expression, while the presence of small naked valleys in other species of the *virilis* group (e.g. D. americana) could be explained by a gain of microRNA expression. The coloured bars represent the spatial expression of each gene in the femur with lighter orange indicating where *sha* expression is post-transcriptionally repressed by *miR-92a.*

Interestingly, we observed that the ectopic trichomes produced by expression of *sha*-ΔUTR in the naked valley are significantly shorter than those on the rest of the leg (S5 Fig). Therefore while *sha* is able to induce trichome formation in these cells, other genes including *CG14395* are also required for their normal morphology. This suggests that GRNs may be able to co-opt regulators, in this case possibly miR-92a, that can act in trans to regulate existing components. Such changes can facilitate phenotypic evolution by phenocopying the effects of ‘hotspot’ genes in contexts where their evolution may be constrained. While trichomes can be lost as a result of changes in *svb* expression, but not *sha* alone, interestingly, over expression of *miR-92a* is also able to suppress trichomes on other structures, including wings [1], presumably through repression of *sha* and other genes like CG14395.

### Other genetic bases for the evolution of leg trichome patterns?

In contrast to larvae, it is unlikely that mutations in *svb* can lead to evolutionary changes in legs to gain trichomes and decrease the size of the naked valley. This is because this gene (and all the other genes necessary for trichome production) is already transcribed in naked cells. In addition, a single *svb* enhancer is able to drive expression throughout the legs including the naked valley. Although other enhancer regions of this gene are able to drive some expression in patches of leg cells, none of these is naked valley-specific. This suggests that evolutionary changes to *svb* enhancers would be unlikely to only affect expression of this gene in the naked valley. It remains possible that binding sites could evolve in this global leg enhancer to increase the Svb concentration specifically in naked valley cells. This could overcome miR-92a-mediated repression of trichomes similar to experiments where *tal* and *ovoB* are over expressed in these cells, or when molecular sponges are used to phenocopy the loss microRNAs [49]. However, this does not seem to have been the preferred evolutionary route in *D. melanogaster* [1] (Fig 5).

Our study also corroborates that *Ubx* represses leg trichomes [42] whereas it promotes larval trichome development through activation of *svb* [46]. Moreover, our results indicate that *Ubx* acts upstream of *miR-92a* in legs because it is unable to repress leg trichomes in the absence of this microRNA. It is possible that *Ubx* even directly activates *miR-92a* since ChIP-chip data indicate that there are Ubx binding sites within the *jigr1*/*miR-92a* locus [50]. Intriguingly, there is no naked valley in *D. virilis*, and *Ubx* does not appear to be expressed in the second legs of this species during trichome development [42] (Fig 5). However naked valleys are evident in other species in the *virilis* and montana groups and it would be interesting to determine if these differences were caused by changes in *Ubx, miR-92a* or even other loci (Fig 5).

### Evolutionary hotspots and developmen*tal* context

To the best of our knowledge, our study is the first to directly compare the expression and function of components of the GRNs underlying formation of similar structures that have evolved in different developmen*tal* contexts. Our results show that the GRNs for trichome production in larval versus leg contexts retain a core set of genes but also exhibit differences in the components used and in their wiring. These differences likely reflect changes that accumulate in GRNs during processes such as co-option [51] and developmen*tal* systems drift [52-54], although it remains possible that the changes have been selected for unknown reasons.

Importantly, we show that the differences in these GRNs may help to explain why they have evolved at different nodes to lead to the gain or loss of trichomes. This supports the suggestion that GRN architecture can influence the pathway of evolution and lead to hotspots for the convergent evolution of traits [17-19,21]. Indeed, such hotspots can also underlie phenotypic changes in different developmen*tal* contexts. For example, *yellow* underlies differences in abdominal pigmentation and wing spot pigmentation among *Drosophila* species [7,11,55,56]. However, we demonstrate that it cannot be assumed that evolutionary hotspots in one development context represent the nodes of evolution in a different context as a consequence of differences in GRN architecture.

Our findings also highlight that the genes that underlie the loss of features might not have the capacity to lead to the gain of the same feature. Therefore, while evolution may be predictable in particular contexts, it is very important to consider developmen*tal* context and whether a trait is lost versus gained. Indeed even when we map the genetic basis of phenotypic change to the causative genes it is important to understand the changes in the context of the wider GRN to fully appreciate how the developmen*tal* program functions and evolves. Since evolution is thought to favour changes with low pleiotropy [19,57-60], the effects of genetic changes underlying phenotypic change should be tested more widely during development. Such an approach recently revealed that *svb* enhancers underlying differences in larval trichomes are actually also used in other contexts [41]. Interestingly, *miR-92a* is employed in several roles, including self-renewal of neuroblasts [44], germline specification [45], and circadian rhythms [61]. It remains to be seen if the changes in this microRNA underlying naked valley differences also have pleiotropic consequences, and therefore if natural variation in naked valley size is actually a pleiotropic outcome of selection on another aspect of *miR-92a* function.

## Materials and Methods

### Fly strains, husbandry and crosses

Fly strains used in this study are listed in S4 Table. Flies were reared on standard food at 25 °C if not otherwise indicated.

Replacement of the P{lacW}l(3)S011041 element, which is inserted 5’ of the *tal* gene, by a P{GaWB} transposable element was carried out by mobilization in *omb-Gal4; +/CyO Δ2–3; l(3)S011041/TM3Sb* flies as described in [30]. Replacements were screened by following UAS-GFP expression in the progeny. The P{GaWB} element is inserted in the same nucleotide position as P{lacW}S011041. Clonal analysis of *tal* S18.1 and *svbR9* alleles were performed as previously described [62].

A transgenic line that contains the cis-regulatory region of *svb* upstream of a GFP reporter (*svb*BAC-GFP) [41] was used to monitor *svb* expression. Legs of pupae were dissected 24 h after puparium formation (hAPF), fixed and stained following the protocol of Halachmi et al. (2012) [63], using a chicken anti-GFP as primary antibody (Aves Labs, 1:250) and an anti-chicken as secondary (AlexaFluor 488, 1:400). Images were obtained on a confocal microscope with a 60X objective. SUM projections of the z-stacks were generated after background subtraction. A filter median implemented in ImageJ software [64] was applied. The proximal femur image was reconstructed from two SUM projections using Adobe Photoshop.

### Measurement of trichome length

For trichome length measurements, T2 legs were dissected, mounted in Hoyer’s medium/lactic acid 1:1 and imaged under a Zeiss Axioplan microscope using ProgRes^®^ MF cool camera (Jenaoptik, Germany). Trichomes on dis*tal* and proximal femurs were measured and analysed using ImageJ software [64]. Statistical analyses were done in R version 3.4.2 [65].

### RNA-Seq

Pupae were collected within 1 hAPF and allowed to develop for another 20 to 28 h at 25 °C. Second legs were dissected in PBS from approximately 80 pupae per replicate and kept in RNAlater. RNA was isolated using phenol-chloroform extraction. This was done in three replicates for two different strains (*e*^4^, *wo*^1^, *ro*^1^ and OregonR). Library preparation and sequencing (75 bp paired end) were carried out by Edinburgh Genomics. Reads were aligned to *D. melanogaster* genome version 6.12 [66] using TopHat 2.1.1. [67]. Transcripts were quantified using Cufflinks 2.2.1 and differential expression analysis conducted using Cuffdiff [68] (S1-7 Files). Genes expressed below or around 1 FPKM were considered not expressed. Raw reads will be deposited in the Gene Expression Omnibus.

### ATAC-seq

Pupae were reared and dissected as described above. Dissected legs were kept in ice cold PBS. Leg cells were lysed in 50 µl Lysis Buffer (10 mM Tris-HCl, pH = 7.5; 10 mM NaCl; 3 mM MgCl2; 0.1 % IGEPAL). Nuclei were collected by centrifugation at 500 g for 5 min. Approximately 60,000 nuclei were suspended in 50 µl Tagmentation Mix [25 µl Buffer (20 mM Tris-CH COO^−^, pH = 7.6; 10 mM MgCl_2_; 20 % Dimethylformamide); 2.5 µl Tn5 Transposase; 22.5 µl H2O] and incubated at 37 °C for 30 min. After addition of 3 µl 2 M NaAC, pH = 5.2 DNA was purified using a QIAGEN MinElute Kit. PCR amplification for library preparation was done for 15 cycles with NEBNext High Fidelity Kit; primers were used according to [69]. This procedure was carried out for three replicates for each of two strains (*e*^4^, *wo*^1^, *ro*^*1*^ and OregonR). Paired end 50 bp sequencing was carried out by the Transcriptome and Genome Analysis Laboratory Göttingen, Germany. Reads were end-to-end aligned to *D. melanogaster* genome version 6.12 (FlyBase) [66] using bowtie2 [70]. After filtering of low quality reads and removal of duplicates using SAMtools [71,72], reads were re-centered according to [69]. Peaks were called with MACS2 [73] and visualisation was done using Sushi [74] (S8 and S9 Files).

## Acknowledgements

We thank Georgina Haines-Woodhouse for technical assistance, Daniel Leite for help with bioinformatics, and members of the McGregor lab and Maike Kittelmann for comments and suggestions throughout the project. We also thank Francois Payre (University of Toulouse), David Stern (Janelia Farm), Fen-Biao Gao (UMass Med School) and the Vienna Drosophila RNAi Center for providing fly stocks. This work was funded by DFG Research Fellowships to SK (Ki 1831/1-1) and FAF (FR 3929/1-1), a BBSRC DTP studentship to ADB, grants from Ministerio de Economía y Competitividad (BFU2016-74961-P) and the Andalusian Government (BIO-396) to JLGS, and an Austrian Science Fund (FWF) Fellowship to APM (M1059-B09). RNA library preparation and sequencing were carried out by Edinburgh Genomics, The University of Edinburgh. Edinburgh Genomics is partly supported through core grants from NERC (R8/H10/56), MRC (MR/K001744/1) and BBSRC (BB/J004243/1).

## Supporting Information

**S1 Fig.**
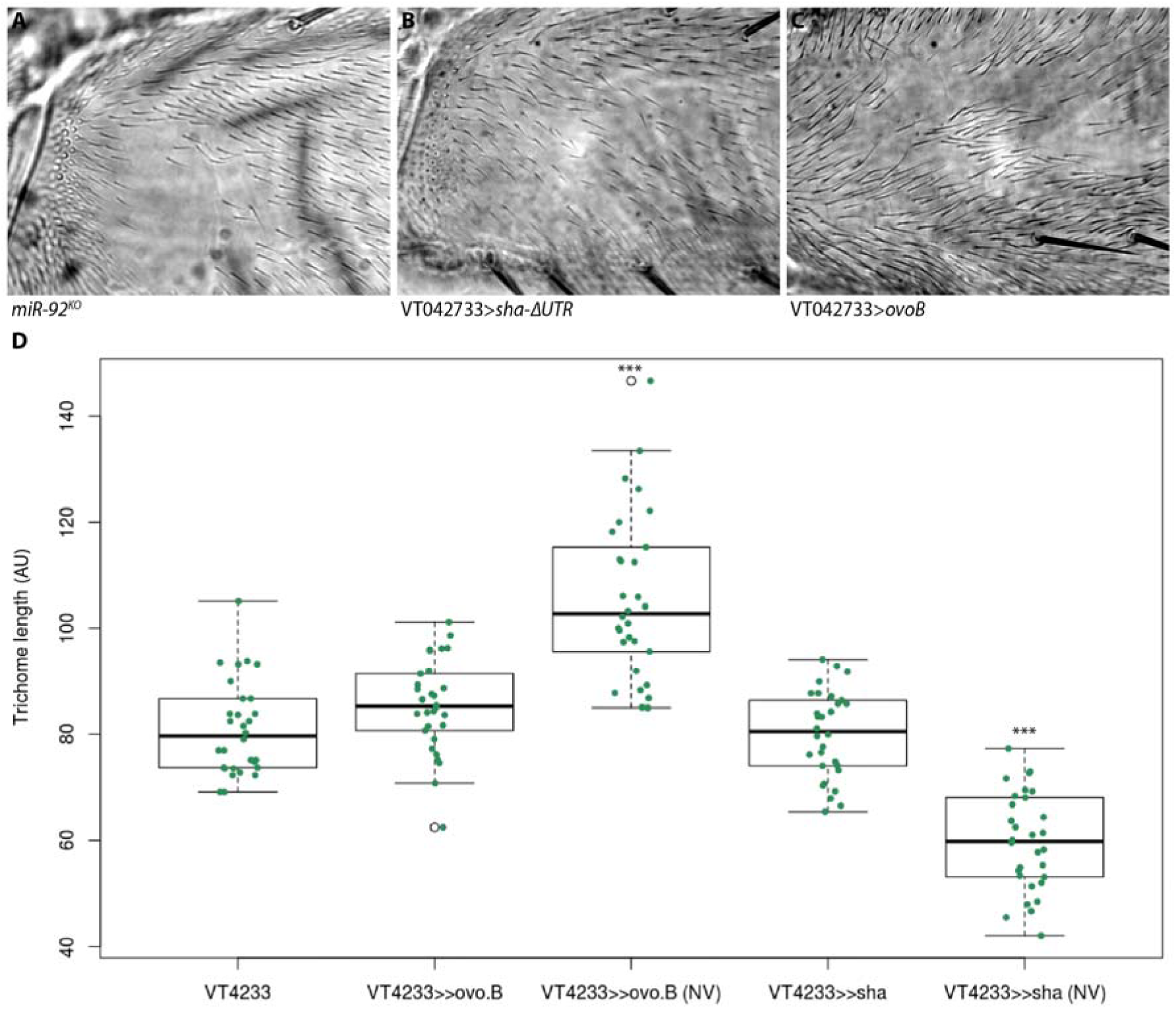
Trichomes gained ectopically in the naked valley have different morphologies. (A) Trichomes gained in the naked valley after loss of *miR-92a* and miR-92b have a similar morphology as trichomes on the more dis*tal* femur. Trichomes gained after ectopic expression of *sha*-ΔUTR (B) are significantly shorter, while trichomes developing after expression of *ovoB* (C) are significantly longer than on the remaining femur. (D) Trichomes on the more dis*tal* femur have a similar length as in the driver line (VT42733) regardless of whether *ovoB* or *sha* are expressed under its control, but trichomes gained in the naked valley are significantly longer or shorter, respectively (p<0.001). Tukey’s multiple comparison test was used to test for significance.

**S2 Fig.**
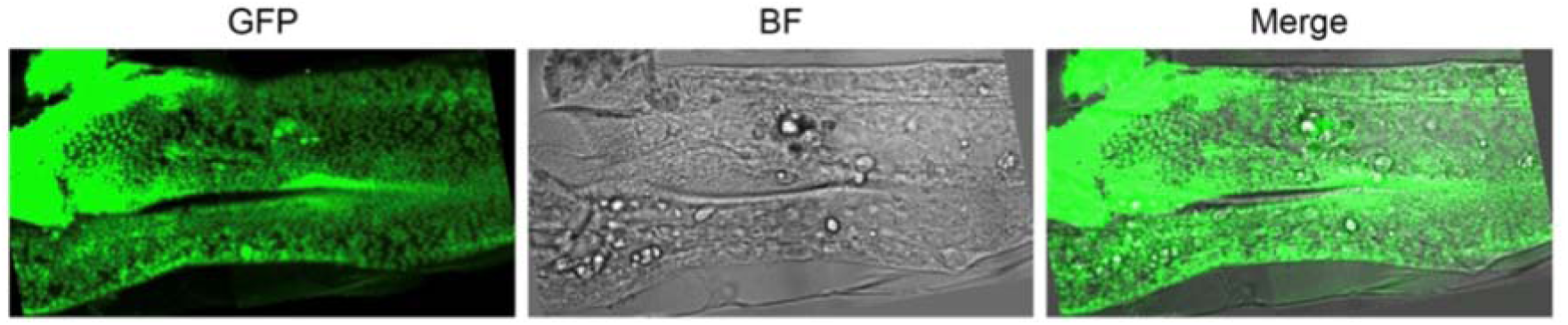
GFP expression driven by *svb*BAC-GFP. GFP is expressed throughout the posterior femur of a T2 leg at 24 hours APF.

**S3 Fig.**
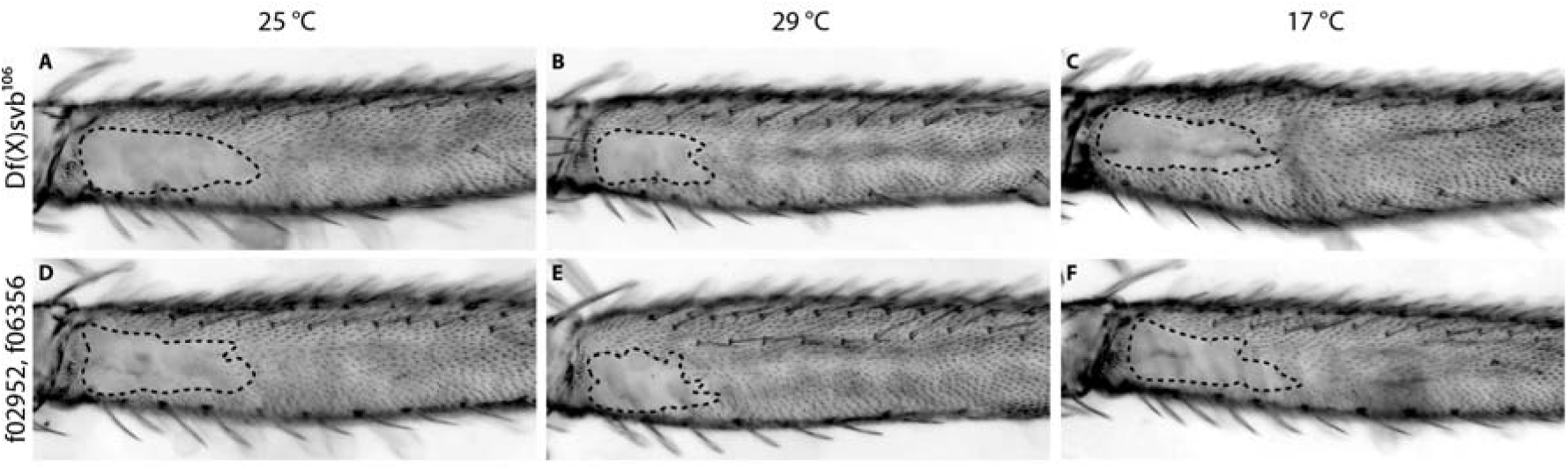
Naked valley size in deficiency line Df(X)106 and control line f02952,f06356. The control line still contains both pBac insertions used to generate the deficiency [5,41]. There is no detectable difference in naked valley size or trichome density between deficiency and control flies at 25 °C, 29 °C, or 17 °C.

**S4 Fig.**
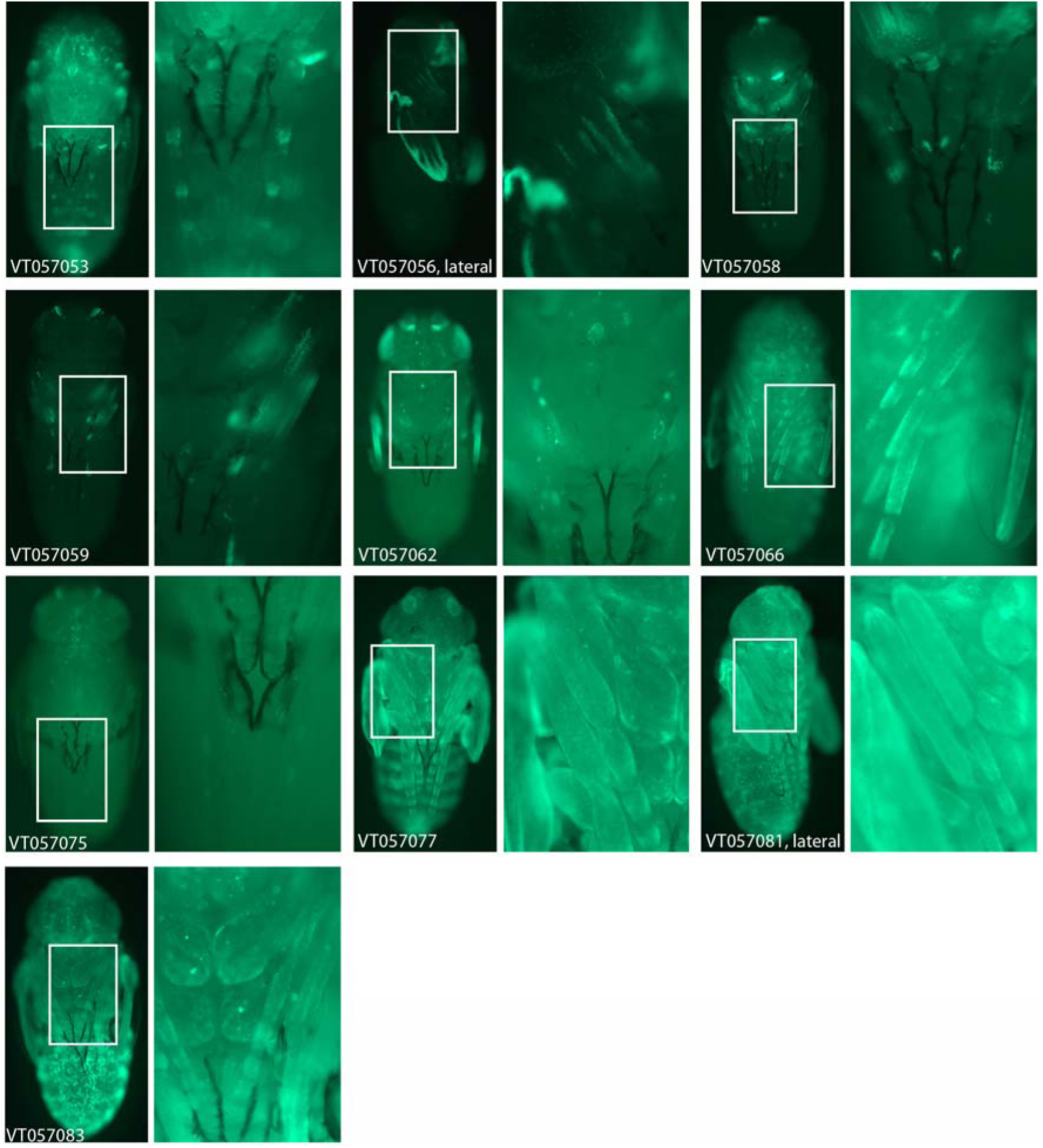
Expression of GFP under control of different VDRC GAL4 drivers in pupae at 22-26 hAPF. All tested drivers show some expression in T2 legs as well as in other pupal tissues.

**S5 Fig.**
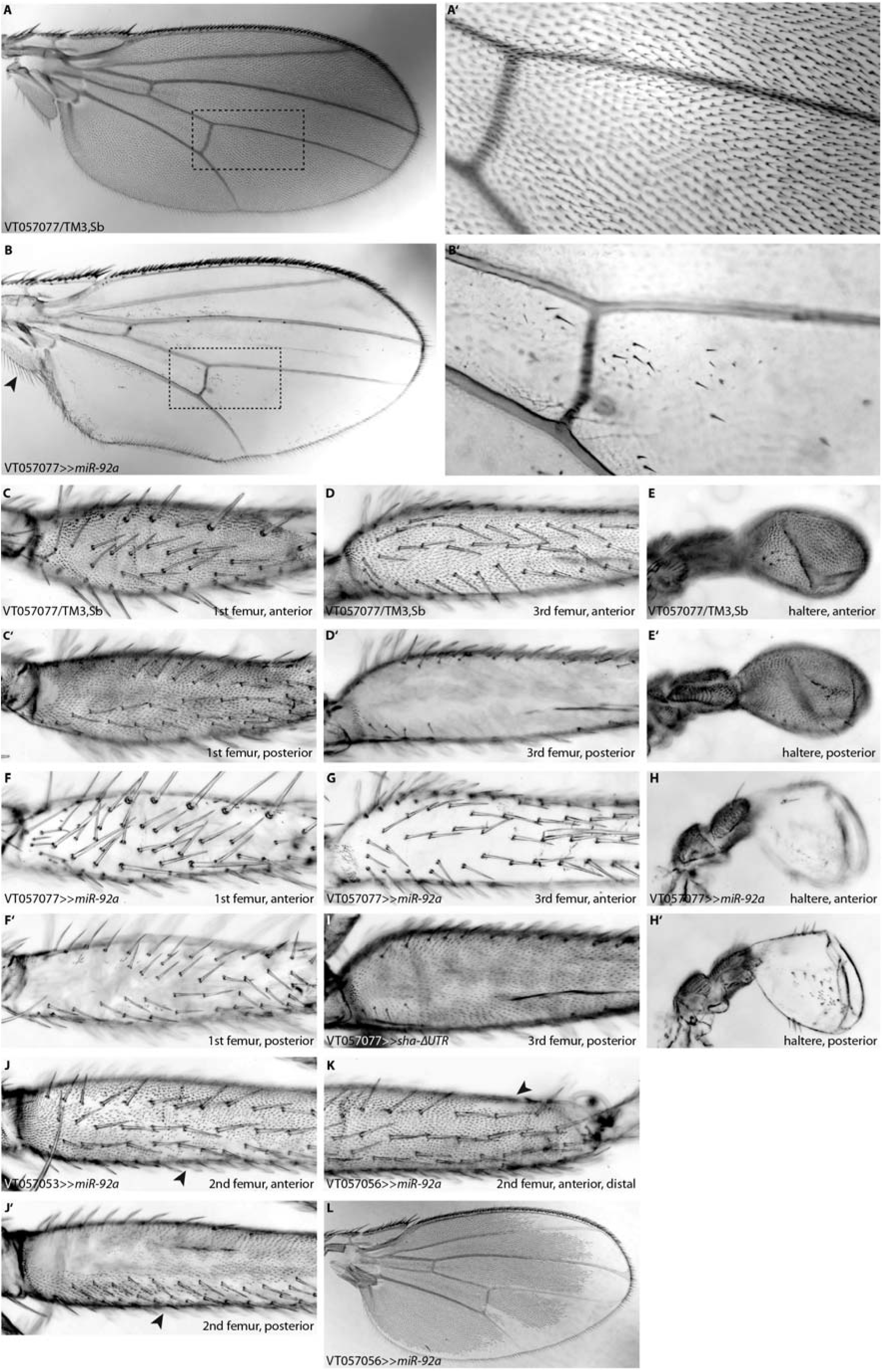
Expression of *miR-92a* and *sha*-ΔUTR under control of different VT Gal4 drivers. (A, A’, B, B’) Trichomes on the wing are largely repressed upon expression of *miR-92a* under control of VT057077. Note that trichomes on the alula (arrowhead in B) develop normally. Also trichomes on T1 and T3 legs (C, C’ D, F, F’, G) and on the halteres (E, E’, H, H’) are repressed when *miR-92a* is driven by VT057077. (I) Driving *sha*-ΔUTR under control of VT057077 leads to ectopic formation of trichomes on the posterior T3 leg (compare to D’). (J, J’) Trichomes on the ventral side of the femur are partially repressed when *miR-92a* is expressed under control of VT057053. Trichomes are repressed in a patch on the dorsal side of the dis*tal* T2 femur (K) and around the rim of the dis*tal* wing (L) after expression of *miR-92a* under control of VT057056.

**S6 Fig.**
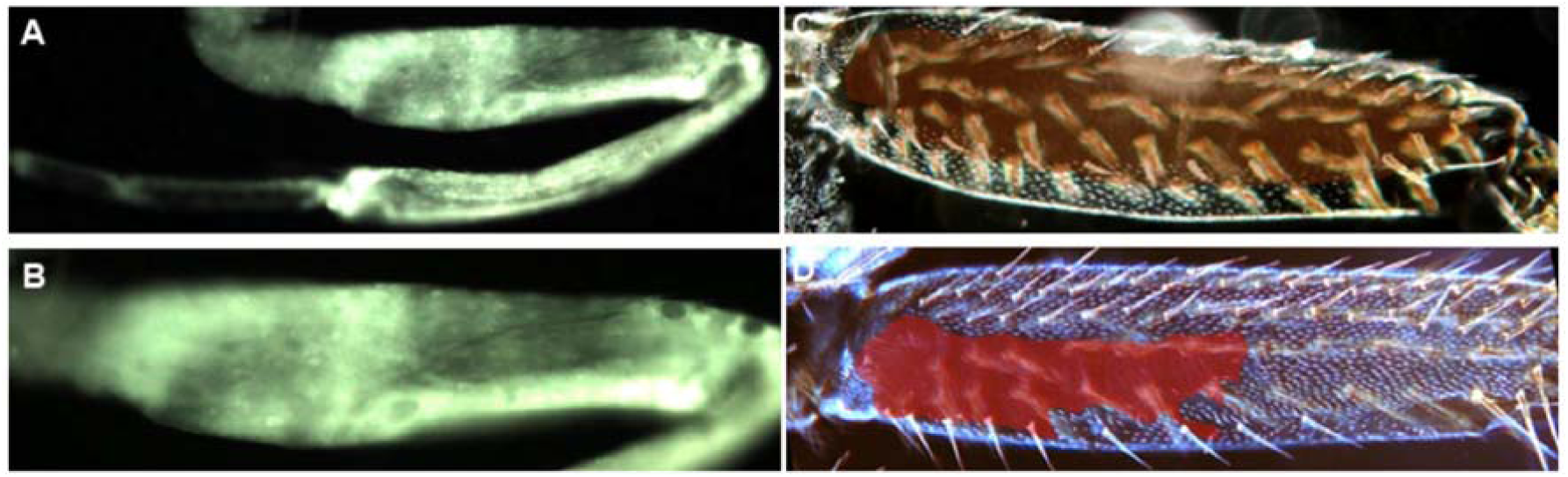
GFP expression driven by *tal*^*lacZ*^ Gal4. GFP is expressed throughout all the leg segments (A) and in the femur (B) of the second leg. Mutant clones of *tal*^*s18*^(C) (brown *sha*ded area) and *svb* ^R9^ (D) (red *sha*ded area) lack trichomes on the femur of a second leg.

**S7 Fig.**
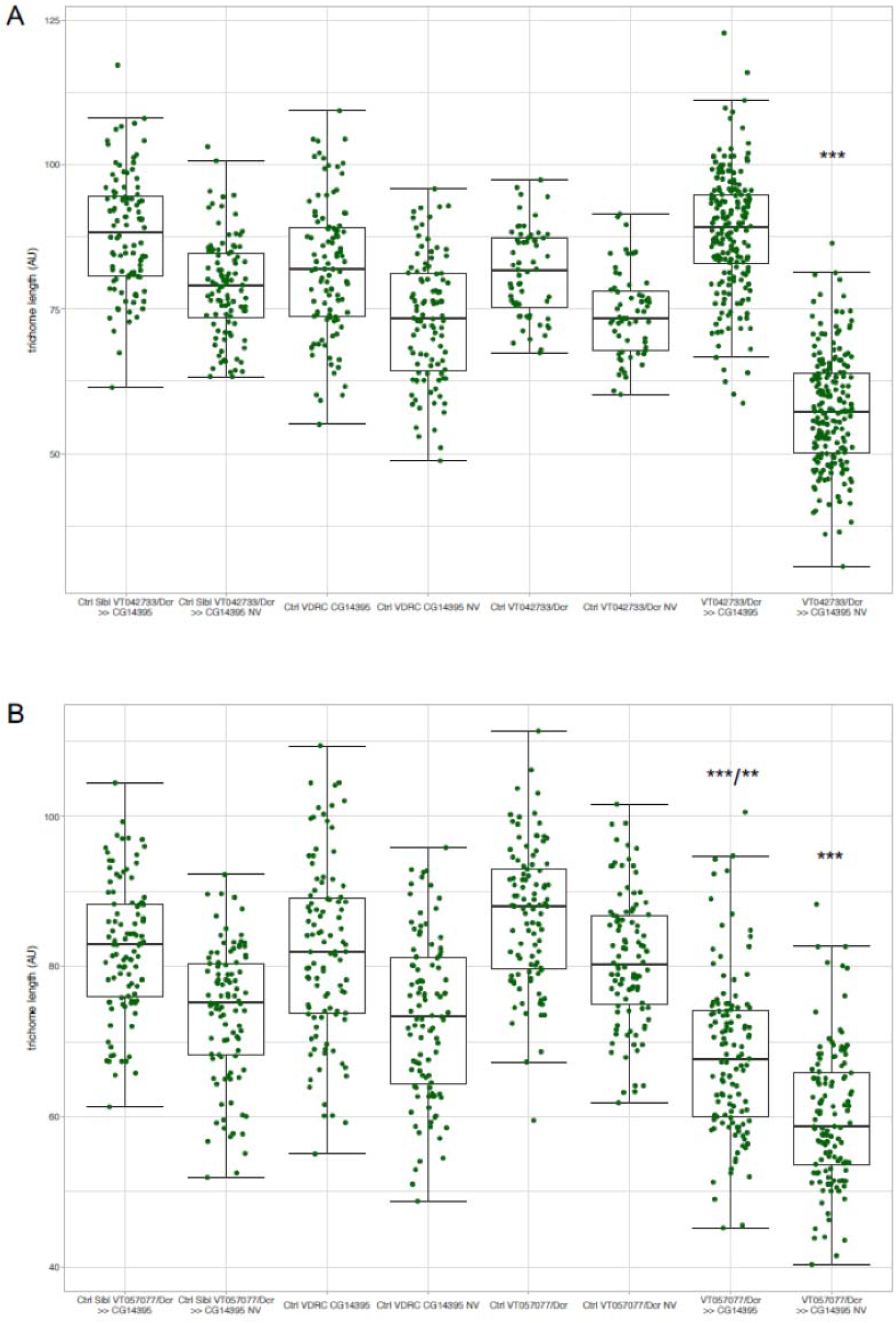
Analysis of trichome length after knockdown of CG14395. Expression of the RNAi construct and *UAS-Dicer* was under control of GAL4 driver lines VT042733 (drives in the proximal femur) and VT057077 (drives in the whole leg). Box plots show the length of trichomes in the dis*tal* part of the posterior femur and around the naked valley (NV). Parents (UAS-Dcr/CyO;VT042733/TM6B or UAS-Dcr/CyO;VT057077/TM6B females, VDRC *CG14395* males) and siblings without knockdown effect were used as controls (Ctrl). (A) Trichomes developing after knockdown of *CG14395* in the proximal femur are significantly shorter around the naked valley area than on the remaining femur (dis*tal* part) and on femurs of the controls (p < 0.001). Data are normally distributed (Shapiro-Wilk normality test). Tukey’s multiple comparison test was used to test for significance. (B) After knockdown of *CG14395* in the whole leg trichomes are significantly shorter both around the naked valley area and on the remaining femur (p < 0.001 and p < 0.01). Note that some controls show significantly different trichome lengths. Data are not normally distributed (Shapiro-Wilk normality test). Kruskal-Wallis and pairwise comparisons using Wilcoxon rank sum test were used to test for significance.

**S1 Table. Comparison of RNA expression levels of upstream trichome network genes for T2 legs at 24 hAPF from two strains with different naked valley size (*e***^**4**^, ***wo***^**1**^, ***ro***^**1**^ **(eworo, large naked valley) and OregonR (OreR, small naked valley)).** Genes are sorted by gene name. Two rows for a gene indicate alternative transcription start sites. Expression level in fragments per kilobase per million (FPKM) with low and high confidence values, base 2 log of the fold change, and p and q values are given after mapping with TopHat 2.1.1, transcriptome assembly with Cufflinks 2.2.1 and Cuffmerge, and comparison with Cuffdiff 2.2.1 [67,68]. q values are false discovery rate (FDR)-corrected p values.

**S2 Table. Comparison of RNA expression levels of genes downstream of *svb* and independent of *svb* [29,32] for T2 legs at 24 hAPF from two strains with different naked valley size [*e***^**4**^, ***wo***^**1**^, ***ro***^**1**^ **(eworo, large naked valley) and OregonR (OreR, small naked valley)]**. Genes are sorted by gene name. Two or more rows for a gene indicate alternative transcription start sites. Expression level in fragments per kilobase per million (FPKM) with low and high confidence values, base 2 log of the fold change, and p and q values are given after mapping with TopHat 2.1.1, transcriptome assembly with Cufflinks 2.2.1 and Cuffmerge, and comparison with Cuffdiff 2.2.1 [67,68].

**S3 Table. Comparison of RNA expression levels of genes independent of *svb* [32,35] for T2 legs at 24 hAPF from two strains with different naked valley size [*e***^**4**^, ***wo***^**1**^, ***ro***^**1**^ **(eworo, large naked valley) and OregonR (OreR, small naked valley)].** Genes are sorted by gene name. Two or more rows for a gene indicate alternative transcription start sites. Expression level in fragments per kilobase per million (FPKM) with low and high confidence values, base 2 log of the fold change, and p and q values are given after mapping with TopHat 2.1.1, transcriptome assembly with Cufflinks 2.2.1 and Cuffmerge, and comparison with Cuffdiff 2.2.1 [69, 70]. q values are FDR-corrected p values.

**S4 Table. Fly strains used.**

**S1 File. FPKM values (with high and low confidence values) after transcript quantification with cufflinks for Oregon R replicate 1.**

**S2 File. FPKM values (with high and low confidence values) after transcript quantification with cufflinks for Oregon R replicate 2.**

**S3 File. FPKM values (with high and low confidence values) after transcript quantification with cufflinks for Oregon R replicate 3.**

**S4 File. FPKM values (with high and low confidence values) after transcript quantification with cufflinks for *e,wo,ro* replicate 1.**

**S5 File. FPKM values (with high and low confidence values) after transcript quantification with cufflinks for *e,wo,ro* replicate 2.**

**S6 File. FPKM values (with high and low confidence values) after transcript quantification with cufflinks for *e,wo,ro* replicate 3.**

**S7 File. FPKM values (with high and low confidence values) for both Oregon R and *e,wo,ro* after comparison with cuffdiff.**

**S8 File. Oregon R *svb* locus ATAC-seq peaks (called with MACS2) with information about position, summit position, height, -log10 (p and q values), and enrichment.**

**S9 File. e,wo,ro *svb* locus ATAC-seq peaks (called with MACS2) with information about position, summit position, height, -log10 (p and q values), and enrichment.**

