## Supplementary Materials for "Gene regulatory network architecture in different developmental contexts influences the genetic basis of morphological evolution"

| **Fly strain** | **Source, stock number (if applicable)** |
| --- | --- |
| e^4^, wo^1^, ro^1^ | Bloomington #496 |
| OregonR | N. Posnien, Goettingen, Germany |
| Df(X)svb^108^ | D. Stern, Janelia Farm |
| e03292, f02952 | D. Stern, Janelia Farm |
| Df(X)svb^106^ | D. Stern, Janelia Farm |
| f02952, f06352 | D. Stern, Janelia Farm |
| VT057050 | VDRC #213086 |
| VT057053/TM3, Sb | VDRC #206968 |
| VT057056 | VDRC #207434 |
| VT057058/TM3, Sb | VDRC #207325 |
| VT057059 | VDRC #206729 |
| VT057062 | VDRC #205634 |
| VT057066 | VDRC #205471 |
| VT057075 | VDRC #207288 |
| VT057077/TM3, Sb | VDRC #205391 |
| VT057081/TM3, Sb | VDRC #206590 |
| VT057083 | VDRC #206605 |
| VT057087 | VDRC #206295 |
| VT042733 | VDRC #214040 |
| UAS-Stinger | Bloomington #65402 |
| UAS-miR-92a | E. Lai (Memorial Sloan Kettering Cancer Center, New York, USA) |
| UAS-sha-ΔUTR | Bloomington #32096 |
| UAS-svb | F. Payre, Toulouse, France |
| UAS-ovoB | F. Payre, Toulouse, France |
| UAS-Ubx/TM3, Ser | Bloomington #911 |
| UAS-tal | J.P. Couso and J.I. Pueyo-Marques |
| UAS-CG14395 RNAi | VDRC #17517 |
| UAS-Dcr/Cyo; VT042733/TM6B | J.P. Couso and J.I. Pueyo-Marques; VDRC #214040 |
| UAS-Dcr/Cyo; VT057077/TM6B | J.P. Couso and J.I. Pueyo-Marques; VDRC #205391 |
| wg-Gal4 | Bloomington #4918 |
| miR-92^KO^ | F.-B. Gao (University of Massachusetts Medical School, Worcester, Massachusetts, USA) |
| svb^PL107^/FM0 | F. Payre, Toulouse, France |
| y, svb^R9^, FRT19A | F. Payre, Toulouse, France |
| tal^S18^, FRT82B | J.P. Couso and J.I. Pueyo-Marques |
